# Cellular heterogeneity and *MTH1* play key roles in galactose mediated signaling of the GAL switch to utilize the disaccharide melibiose

**DOI:** 10.1101/2021.06.16.448739

**Authors:** Rajesh Kumar Kar, Paike Jayadeva Bhat

**Author notes:** For further correspondence or. National Institute of Dental and Craniofacial Research, National Institutes of Health, Bethesda, Maryland-20892 (Rajesh Kumar Kar).

## Abstract

Yeast metabolizes the disaccharide melibiose by hydrolyzing it into equimolar concentrations of glucose and galactose by *MEL1*-encoded α-galactosidase. Galactose metabolizing genes (including *MEL1*) are induced by galactose and repressed by glucose, which are the products of melibiose hydrolysis. Therefore, how melibiose catabolization and utilization take place by circumventing the glucose repression is an enigma. Other than the galactose metabolizing genes *MTH1*, a negative regulator of glucose signal pathway has Gal4p binding sites and is induced by galactose and repressed by high glucose concentration. But, at low or no glucose *MTH1* along with its paralogue *STD1* represses hexose transporters, that are involved in glucose transport. This sort of tuning of glucose and galactose regulation motivated us to delineate the role of *MTH1* as a regulator of *MEL1* expression and melibiose utilization. The deletion mutant of *MTH1* shows growth defect on melibiose and this growth defect is enhanced upon the deletion of both *MTH1* and its paralogue *STD1*. Microscopy and flowcytometry analysis, suggest, that even though *MEL1* and *GAL1* promoter are under Gal4p and Gal80p regulation, upon deletion of *MTH1* it hampers only *MEL1* expression, but not the *GAL1* gene expression. By using 2-Deoxy galactose toxicity assay, we observed phenotypic heterogeneity in cells grown on melibiose i.e. after cleaving of melibiose a fraction of cell population utilizes glucose and another fraction utilizes galactose and coexist together. Understanding *GAL/MEL* gene expression patterns in melibiose will have great implication to understand various other complex sugar utilizations, tunable gene expressions and complex feedback gene regulations.

**Significance:** Sugar metabolism is an important phenomenon to understand the regulation of gene expression. Glucose is the most preferred carbon source. Yeast follows glycolytic pathways like cancer cells for metabolism of sugars and understanding this will throw more light to the metabolism of cancer cells. In this communication we observed cell-to-cell heterogeneity in yeast cells playing a key role in metabolism of a complex disaccharide melibiose, which gets cleaved into glucose and galactose by α-galactosidase. Glucose represses α-galactosidase and galactose induces it. Because of the heterogeneous population of cells one fraction consumes glucose liberated by melibiose hydrolysis, therefore it is not sufficient to repress α-galactosidase and other GAL genes. Therefore, GAL genes are expressed and help in metabolizing melibiose and galactose.

## Introduction

Of all the sugars, glucose is the most preferred carbon source and glucose induces some genes and represses many other genes (Johnston, 1999; Lafuente et al., 2000). For example, the disaccharide sucrose is hydrolyzed into glucose and fructose by invertase, the trisaccharide raffinose is hydrolyzed into fructose and melibiose by invertase and the disaccharide melibiose is hydrolyzed into glucose and galactose by α-galactosidase. Therefore, now the question arises, how these varieties of sugars such as di and trisaccharides are utilized in nature by yeast when in all the above cases equimolar, glucose moiety is generated, which is the repressing carbon source for all the genes whose gene product cleaves the complex sugars like sucrose, raffinose or melibiose.

The α-galactosidase encoded by gene *MEL1* is not present in smaller yeasts like *Saccharomyces cerevisiae*, but is present in larger yeasts like *Saccharomyces bayanus, Saccharomyces carlsbergensis* (*S. pastorianus*) etc (Turakainen et al., 1993). The *MEL1* gene has been introduced into *S. cerevisiae* to study various aspects of the *GAL/MEL* genetic switch and used as a reporter for measurement of α-galactosidase expression. It is known that when yeast is grown in galactose as a sole carbon source, the *GAL* genes are actively transcribed (Bhat and Iyer, 2009; Bhat and Murthy, 2001; Johnston, 1987; Lohr et al., 1995). The enzymes coding for galactose utilization are repressed in the presence of glucose by a cascade of molecular mechanisms, and this phenomenon is known as carbon catabolite repression (Carlson, 1999; Johnston, 1999). As discussed earlier, glucose is one of the products of melibiose hydrolysis catalyzed by the *MEL1* gene product α-galactosidase, which by itself is glucose repressed **(Figure S1A)**. Therefore, the molecular mechanism of how α-galactosidase circumvents the glucose repression and cleaves melibiose into equimolar glucose and galactose is still an enigma. A clue to the molecular mechanism of how the *MEL1* gene product α-galactosidase circumvents the glucose repression and is induced by galactose for the hydrolysis of the disaccharide melibiose came from the regulation pattern of *MTH1* (<u>M</u>SN <u>T</u>hree <u>H</u>omologue) **(Figure S1B)**.

Genome-wide location of DNA binding protein analysis suggests that, other than the previously identified galactose metabolizing genes, *MTH1, PCL10, GCY1* and *FUR4* are also regulated by Gal3p-Gal80p-Gal4p interaction (Ren et al., 2000). Among these only *MTH1* is upregulated by galactose (Ren et al., 2000) and is a negative regulator of the *HXT* (Hexose transporters) genes, which are involved in the glucose-sensing signal transduction pathway **(Figure S1B)** (Hubbard et al., 1994; Lafuente et al., 2000), but why they are under Gal4p regulation and its role in *GAL*/*MEL* pathway is not clearly understood (Ren et al., 2000). The Mth1p represses the *HXT* genes by physically binding to Rgt1p, along with its paralogue Std1p (<u>S</u>uppressor of <u>T</u>ATA binding protein <u>D</u>eletion), and recruits the general co-repressors Ssn6p and Tup1p in the absence of glucose or at very low glucose concentration (Kim et al., 2003; Malave and Dent, 2006; Ozcan et al., 1996; Smith and Johnson, 2000) **(Figure S1B)**. These general co-repressors Ssn6p and Tup1p recruit a large number of global repressors to inhibit the transcription of *HXT* genes along with the main repressor Rgt1p (Malave and Dent, 2006; Smith and Johnson, 2000). The Mth1p has been shown to work as an adaptor to bring Ssn6p and Tup1p to Rgt1p and modulates the PKA (Protein kinase A) dependent phosphorylation of Rgt1p (Roy et al., 2013). Mth1p also has been shown to bind to the C-terminal tail of glucose sensors Rgt2p/Snf3p in the presence, as well as absence, of glucose (Lafuente et al., 2000). The glucose sensors Snf3p and Rgt2p are coupled with Casein kinase (Yck1p), which phosphorylates both Mth1p and Std1p in the presence of glucose and degrades it (Moriya and Johnston, 2004).

*MTH1* and *STD1* arose due to whole genome duplication events from its common ancestors (Wolfe and Shields, 1997) and are also shown to have 61% identity with each other (Hubbard et al., 1994). In the ancestor *K. lactis* a single gene *SMS1* (<u>S</u>imilar to *<u>M</u>TH1* and *<u>S</u>TD1*) was identified, which encodes a protein shown to have 52% and 48% identity with Mth1p and Std1p respectively, of *S. cerevisiae* (Hnatova et al., 2008). *MTH1* and *STD1* have also been shown to bind with a different affinity to the C-terminal tail of Snf3p as well as Rgt2p (Schmidt et al., 1999). Glucose stimulates the proteasomal degradation of Mth1p and Std1p (Kim et al., 2006; Moriya and Johnston, 2004; Pasula et al., 2007; Spielewoy et al., 2004), but it induces *STD1* expression and represses *MTH1* expression (Kaniak et al., 2004; Kim et al., 2006), therefore are not redundant, but may have evolved for better regulation of complex sugar utilization. Therefore, we investigated the role of *MTH1*, along with its paralogue *STD1*, for the regulation of tuning of glucose and galactose signaling, which plays a critical role in *MEL1* gene expression that leads to hydrolysis of melibiose.

By using a 2-Deoxy galactose toxicity plate assay we observed cell-to-cell heterogeneity in populations of cells grown on melibiose. One cell population utilizes glucose and the other utilizes galactose i.e. the population consists of cells with physiologically distinct states: one that can utilize glucose with *GAL* genes repressed and the other with *GAL* genes ON, which can utilize galactose. Our microscopy and flowcytometry of melibiose grown cells integrated with *P_GAL1GFP_* and *P_MEL1_mCherry*, suggest that even though both the *GAL1* and *MEL1* promoters are regulated by the same galactose signal-driven Gal3p mediated Gal4p-Gal80p interaction mechanism, Mth1p regulates only the *MEL1*, which has a single Gal4p binding site, but not *GAL1* promoter, which has two Gal4p binding sites. Mth1p regulates *MEL1* for the expression of α-galactosidase by some unknown interaction other than the galactose signal-driven Gal3p-mediated Gal4p-Gal80p interaction mechanism and for balancing the tuning of glucose and galactose concentration upon hydrolysis of the disaccharide melibiose.

The implications of understanding this mechanism will help to understand more complex patterns of gene regulation in other complex sugars, like sucrose and raffinose utilization, but also to understand the reversible conversion of ethanol to acetaldehyde by *ADH1* and *ADH2* genes in the yeast, which are controlled by feedback mechanisms.

## Results

### Melibiose grown cells show cellular heterogeneity

To understand the complex *GAL/MEL* gene expression behavior of a population of cells utilizing melibiose as a sole carbon source, the following assay was designed. To determine the fraction of cells that utilize glucose and galactose upon hydrolysis of melibiose, the melibiose and galactose pre-grown cells were spotted onto glycerol/lactate (gly/lac) and gly/lac + 0.3% 2-Deoxy-galactose (2-DG) plates **(Figure S2)**. Glycerol is a non-inducible and non-repressible media for *GAL* gene expression. The *GAL* gene expression phenotype of the melibiose and galactose pre-grown cells were monitored in 2-DG plates. If the *GAL* genes are expressed the cells will utilize 2-DG and will be killed and if *GAL* genes are not expressed the cells grow in 2-DG plates. Upon growth in the 2-DG plates, it was observed that all the galactose-grown cells were killed, but in the case of melibiose grown cells only a fraction of cells were killed **(Figure 1A)**. This fraction of the cells that were killed was determined by counting the cells from 2-DG plates. Approximately 50 % cells were killed in 2-DG plates, when pre-grown on melibiose **(Figure 1B)**.

**Figure 1:**
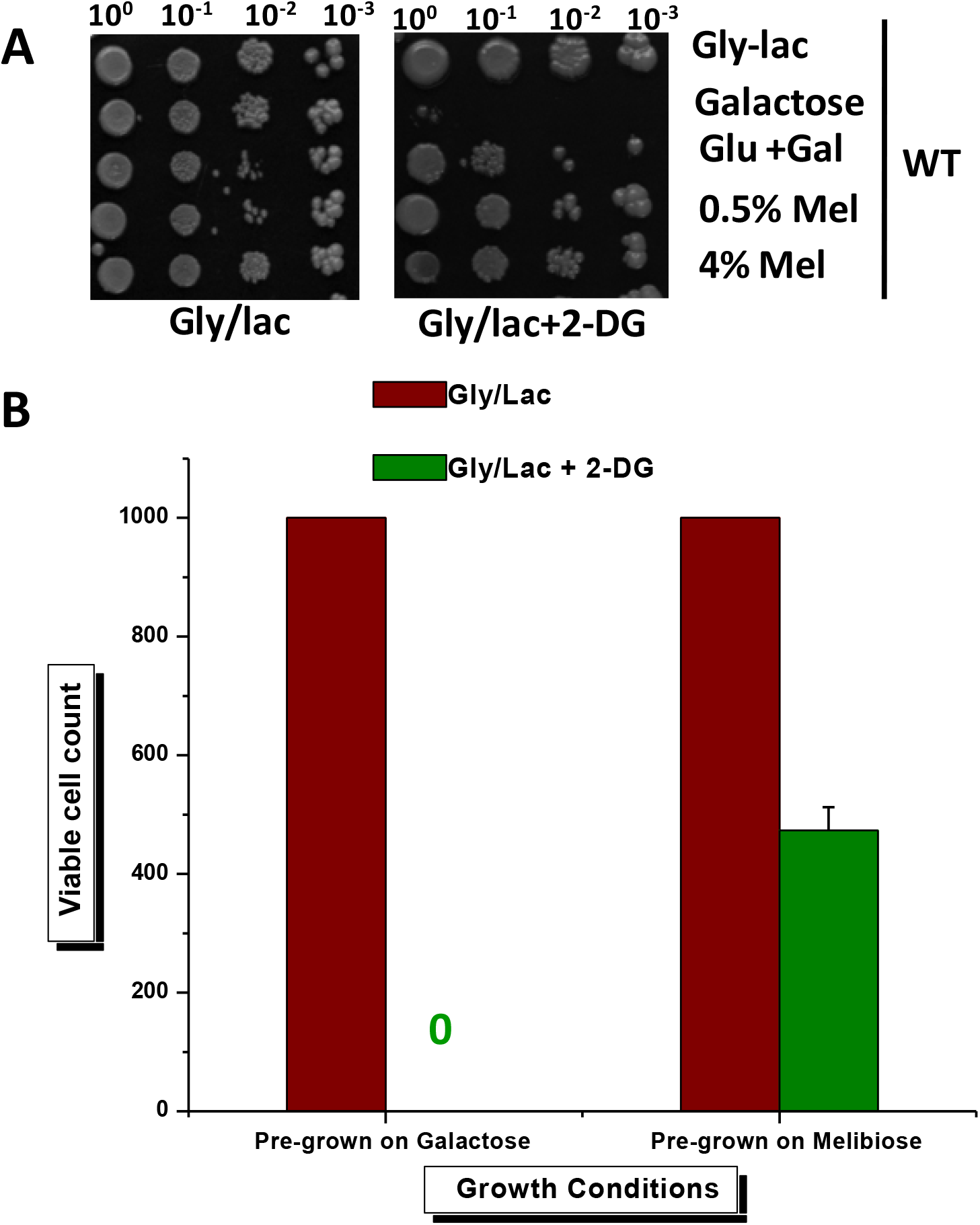
Phenotypic and quantitative analysis of the WT strains pre-grown on the indicated media by 2-DG toxicity assay. **(A)** WT cells pre-grown on the indicated media were serially diluted and spotted onto gly/lac and gly/lac + 2-DG plates. The growth of cells was monitored after 4 days of incubation. **(B)** Equal amount of WT cells pre-grown on galactose or melibiose were plated onto gly/lac and gly/lac + 2-DG plates. The viable cells in 2-DG plates were determined and plotted per 1,000 cells. Red bar represents average cell count of gly/lac plate and green bar represents average cell count of gly/lac + 2-DG plates. The 2-DG plate experiment was repeated thrice, and average and standard deviation are plotted.

### Yeast cells grown on melibiose do not show diauxic shift

Previously it was known that when yeast cells grow on a mixture of sugars like glucose and galactose, glucose is utilized first (Johnston, 1999; Vaulont et al., 2000). The cells pass through a lag phase and only then other carbon sources are utilized. This shift is called the diauxic shift (Monod, 1949; Perez-Samper et al., 2018; Zaman et al., 2008). When the disaccharide melibiose is utilized as the carbon source, it is cleaved by α-galactosidase into equimolar concentrations of glucose and galactose. To demonstrate whether any diauxic shift occurs, when melibiose is utilized as a carbon source, cells were grown in melibiose and growth was monitored by measuring OD at regular intervals. Along with this mixture of glucose + galactose, only glucose or only galactose were used as controls. From the growth experiment we observed that cells grown on the glucose and galactose mixture show a diauxic shift indicated with a blue arrow, which was not observed in melibiose-grown cells **(Figure 2A)**. The red arrow indicates the galactose to ethanol diauxic shift and the green arrow indicates the glucose to ethanol diauxic shift (Peng et al., 2015). The diauxic shift was also confirmed by microscopy **(Figure 2B)** and flowcytometry analysis **(Figure 2C, D** and **E)**. For this, the wild-type (WT) cells integrated with *P_GAL1_GFP* and *P_MEL1_mCherry* were grown on an equimolar (0.25% each) mixture of glucose and galactose. Cells were collected at log phase as well as at stationary phase and the expression of GFP and mCherry were analyzed by microscopy and flowcytometry. GFP and mCherry were analyzed by using FITC and PE-Texas red filters, respectively. At the log phase there was no *P_GAL1_GFP* and *P_MEL1_mCherry* expression, but at the stationary phase both *P_GAL1_GFP* and *P_MEL1_mCherry* expression was observed in all the cells **(Figure 2)**. On the other hand, *P_GAL1_GFP* and *P_MEL1_mCherry* expression was clearly observed in the melibiose grown cells at the log phase **(Figure 4A** and **Figure 5)**.

**Figure 2:**
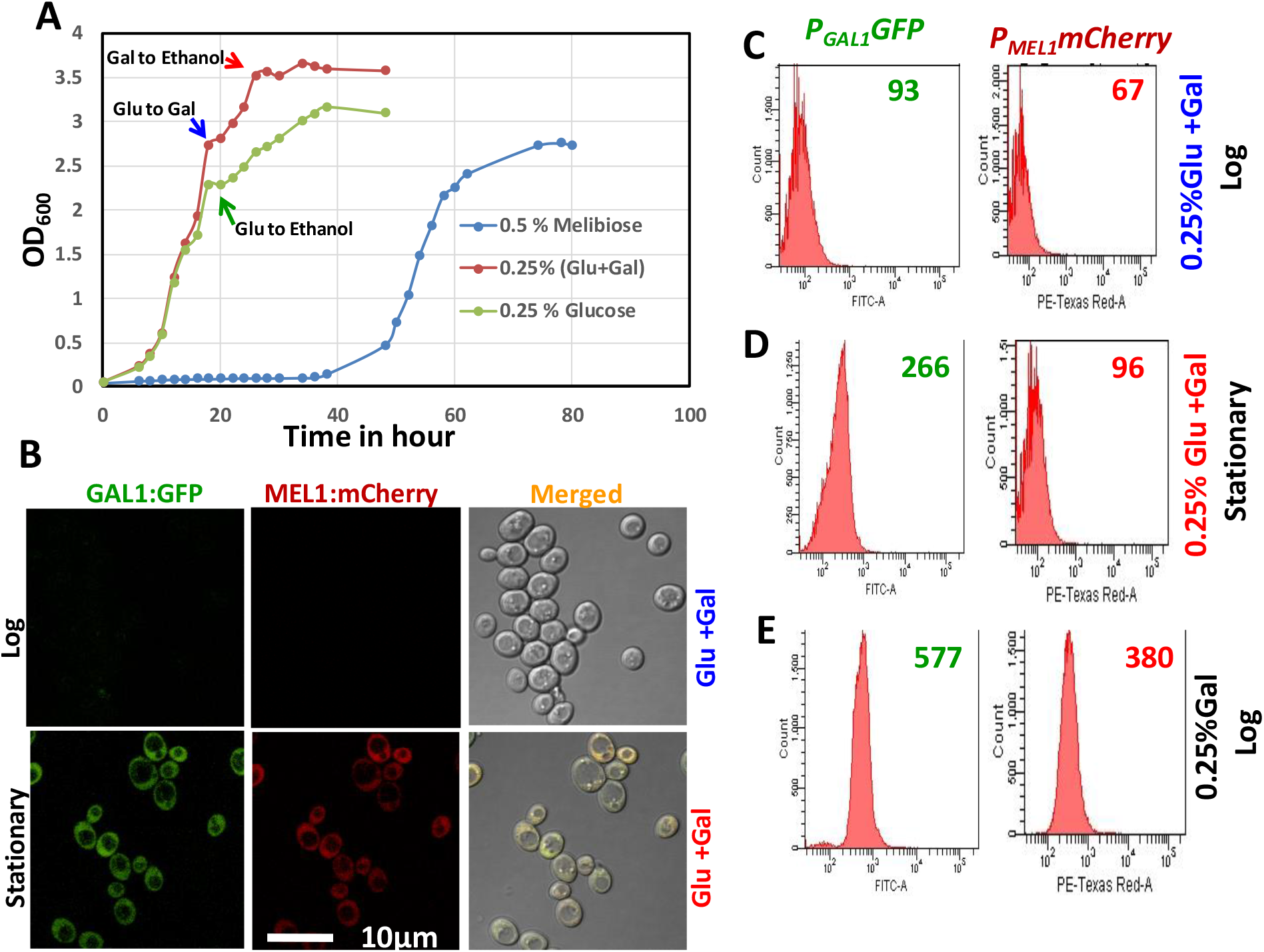
Comparison of the growth phenotypes and gene expression pattern of WT cells in glucose, galactose, melibiose as well as in glucose and galactose mixture. **(A)** Growth curve analysis of WT cells grown on the indicated carbon sources. WT cells were grown on mixture of glucose (Glu) and galactose (Gal) to monitor the diauxic shift (shown by the blue arrow mark). Melibiose and glucose were also taken as the controls to monitor the diauxic shift. Cell growth was monitored by taking OD of the samples collected at 2 hours and few time points at 4- or 6-hours interval as indicated in figure. The red arrow indicates the galactose to ethanol diauxic shift and green arrow indicates the glucose to ethanol diauxic shift. The growth experiment was repeated at least thrice, and average of three biological duplicates are plotted here. **(B)** Microscopic analysis of WT cells integrated with *P_GAL1_GFP* and *P_MEL1_mCherry* grown in the mixture of glucose and galactose. WT cells grown on mixture of glucose and galactose were harvested at log and stationary phase. The GFP and mCherry expression were observed under confocal microscope. Scale bar is 10 μm. flowcytometry analysis of WT cells integrated with *P_GAL1_GFP* and *_PMEL1_mCherry* grown in the mixture of glucose and galactose. Cells grown on glucose and galactose mixture were harvested at **(C)** log and **(D)** stationary phase of the growth and the *P_GAL1_GFP* and *P_MEL1_mCherry* expression were quantified by flowcytometry analysis. **(E)** In this panel, galactose-grown cells are taken as a positive control. The numbers shown in the figure are the mean of fluorescence intensity of either the GFP or mCherry of 50,000 cells. GFP and m-Cherry were measured by using FITC and PE-Texas Red-A filters respectively. The experiment has been performed at least thrice and the pattern was same, therefore only one set of data is presented.

### *MTH1* deletion suppresses the growth on melibiose

*MTH1* and *STD1* are paralogues but only *MTH1* is under Gal4p regulation and represses glucose transport. Therefore, the role of *MTH1* was studied for the growth in melibiose. The *SMS1* gene of *K. lactis* is a homologue of *MTH1* and *STD1*. As *K. lactis* genome has not undergone whole genome duplication, it is known that only *SMS1*, the homologue of *MTH1* and *STD1*, works as a repressor of glucose signaling pathway in *K. lactis*. Therefore, the role of *SMS1* was also studied for the disaccharide lactose growth, which hydrolyzes into equimolar concentrations of glucose and galactose by β-galactosidase. To understand the role of *SMS1*, the WT and *sms1*Δ cells pre-grown in glycerol was spotted onto lactose plate. No growth difference between the WT and *sms1*Δ cells were observed in the lactose, suggesting that *SMS1* may not have very important role in utilization of the disaccharide lactose **(Figure 3A)**. However, a clear growth defect was observed in the disaccharide melibiose in the case of the *mth1*Δ strain but not in the *std1*Δ strain **(Figure 3B)**. Surprisingly the growth defect was enhanced in case of *mth1*Δ*std1*Δ double deletion strain **(Figure 3B)**. These growth defects were also corroborated by the growth curve analysis in two different concentration of melibiose as indicated (**Figure 3C** and **D**).

**Figure 3:**
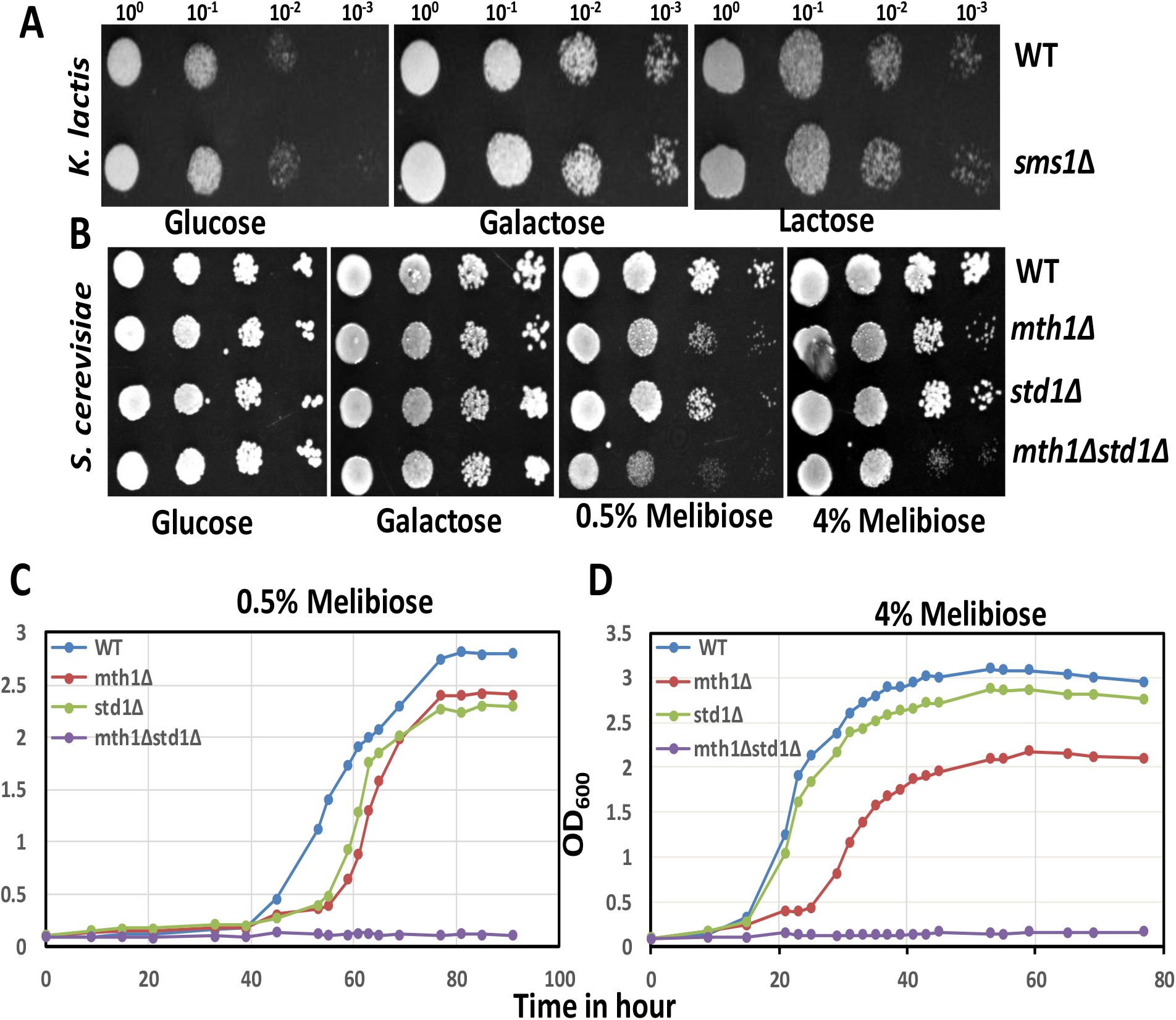
Comparison of growth phenotype of *K. lactis* and *S. cerevisiae* on the disaccharides lactose and melibiose, respectively. **(A)** The growth phenotypic analysis of WT and *sms1*Δ strains of *K*. *lactis* cells on lactose. WT and *sms1*Δ *K. lactis* cells pre-grown on gly/lac were spotted onto the indicated plates. **(B)** Phenotypic analysis of the indicated *S. cerevisiae* strains on melibiose. WT, *mth1*Δ, *std1*Δ and *mth1*Δ*std1*Δ *S. cerevisiae* cells pre-grown on gly/lac were spotted onto the indicated plates. In both *K. lactis* and *S. cerevisiae* growth was monitored after 4 days of incubation. Growth curve analysis of WT, *mth1*Δ, *std1*Δ and *mth1*Δ*std1*Δ strain grown in **(C)** 0.5% and **(D**) 4% of melibiose. The indicated yeast cells were grown on the 0.5% melibiose and 4% melibiose and the growth was monitored by taking OD of the collected samples at the lag phase at 6 hours time interval and at the log phase at 2 hours time interval. The growth experiment was repeated at least thrice, and average of three biological duplicates are plotted here.

### Mth1p regulates *MEL1* but not *GAL1* promoter

To obtain more insights into the *mth1*Δ growth defect phenotype at the molecular level, the *P_GAL1_GFP* and *_PMEL1_mCherry* were integrated into the WT, *mth1*Δ, *std1*Δ and *mth1*Δ*std1*Δ strains. As the growth defect was observed in *mth1*Δ strain and *mth1*Δ*std1*Δ, the *GAL1* and *MEL1* promoter expression were determined by growing the *P_GAL1_GFP* and *P_MEL1_mCherry* integrated cells in galactose and melibiose. From microscopy analysis we observed only a decrease in *MEL1* promoter expression in *mth1*Δ **(Figure 4B)**and *mth1*Δ*std1*Δ **(Figure 4D)**. No change in *MEL1* promoter expression was observed either in WT **(Figure 4A)** or in *std1*Δ **(Figure 4C)** strains and this observation was corroborated by flowcytometry **(Figure 5 and S3)**.

**Figure 4:**
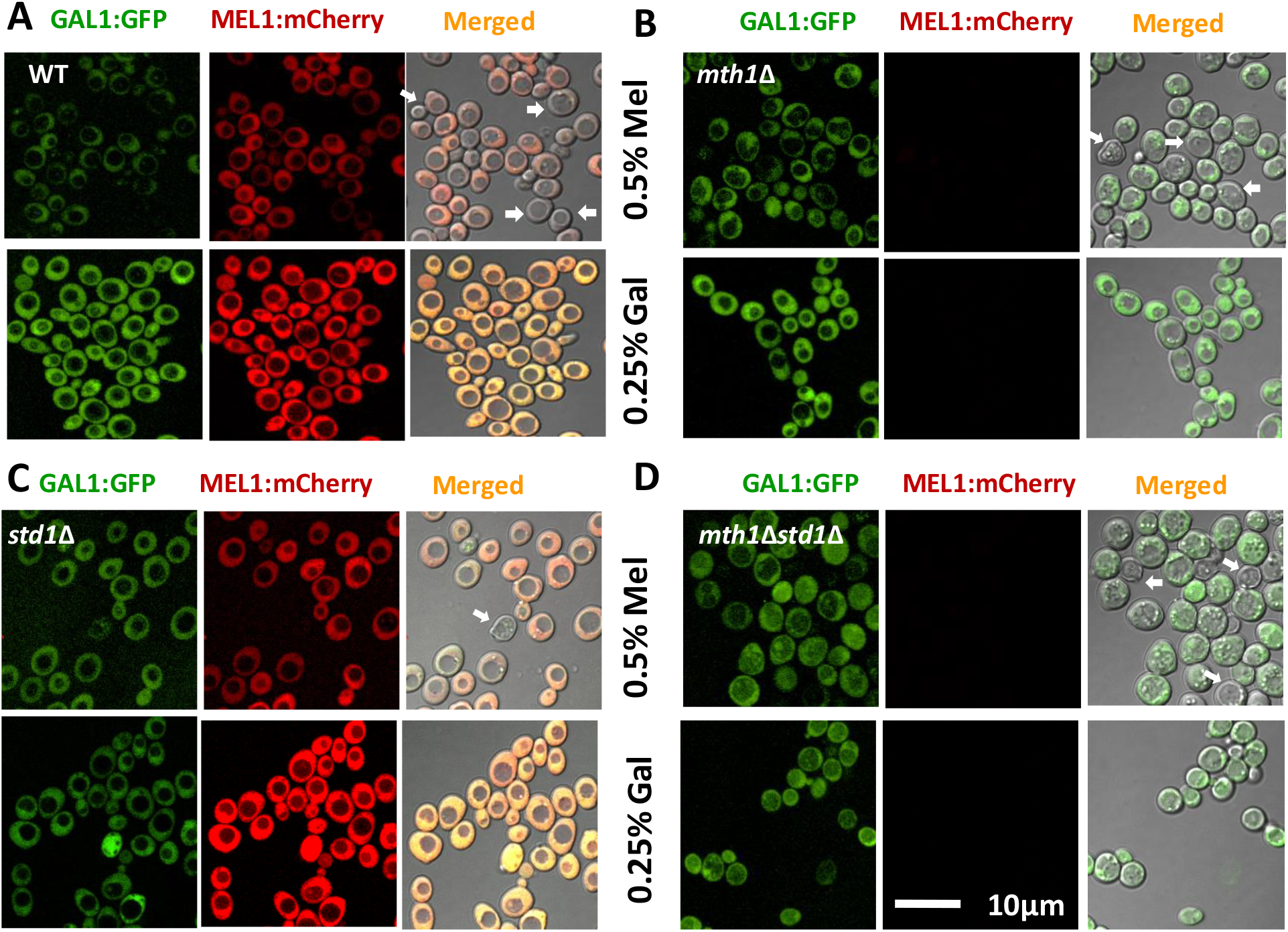
Microscopic analysis of the indicated strains integrated with *P_GAL1_GFP* and *P_MEL1_mCherry* grown in galactose and melibiose. (**A**)WT **(B)** *mth1*Δ, **(C)***std1*Δ and **(D)** *mth1*Δ*std1*Δ cells integrated with *P_GAL1_GFP* and *P_MEL1_mCherey* were grown on the indicated media and monitored in confocal microscope. The scale bar is 10 μm. GFP and m-Cherry were monitored by using EGFP and PE-Texas Red filters respectively. In the population of cells some cells have very high expression of *P_GAL1_GFP* and *P_MEL1_mCherry*, and some cells have very low expression, which are marked with white arrows.

**Figure 5:**
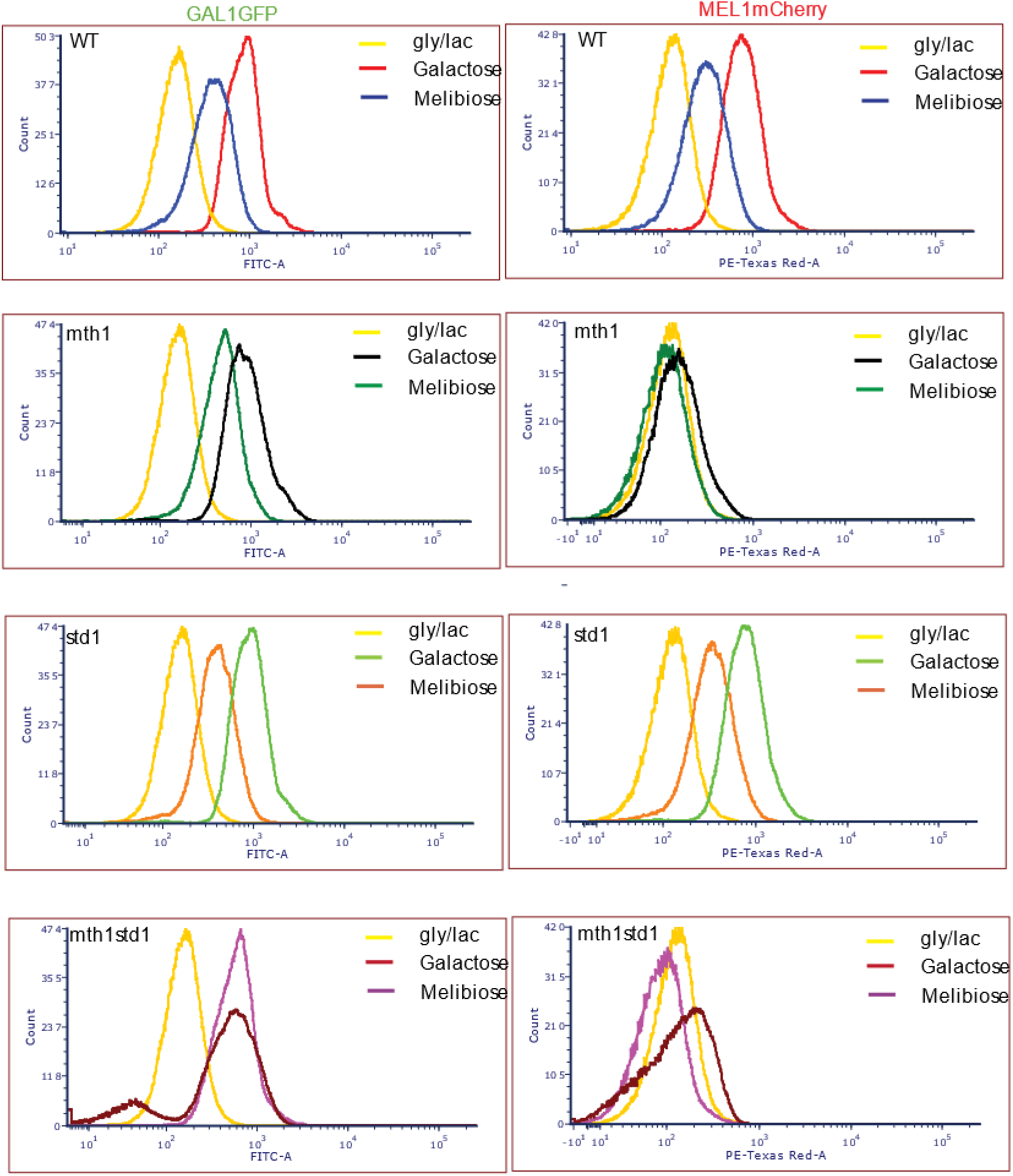
Flowcytometry analysis of the indicated strains integrated with *P_GAL1_GFP* and *P_MEL1_mCherry* grown in galactose and melibiose. WT, *mth1*Δ, *std1*Δ and *mth1*Δ*std1*Δ cells integrated with *P_GAL1_GFP* and *P_MEL1_mCherry* were monitored in flowcytometry after growing in the indicated media and mean fluorescence intensity was plotted. The experiment has been performed at least thrice and the pattern was same, therefore only one set of data is presented. Each data point represents the mean fluorescence intensity of 50,000 cells. GFP and m-Cherry were measured by using FITC and PE-Texas Red-A filters respectively.

Interestingly we observed cell-to-cell heterogeneity in the cells grown on melibiose. In the population of cells, some have very high expression of *P_GAL1_GFP* and *P_MEL1_mCherry*, and some have very low expression, which are marked with white arrows **(Figure 4A)**. But clearly two distinct states of two populations is observed in *mth1*Δ*std1*Δ strains by flowcytometry analysis, where there are two clear peaks **(Figure 5D)**.

### Mth1p does not regulate *MEL1* promoter expression downstream of Gal80p-Gal4p interaction

Since *GAL1* and *MEL1* promoter expression is under Gal4p and Gal80p regulation, Mth1p might be releasing the Gal80p repression from Gal4p in the case of the *MEL1* promoter. To delineate the mechanism of action of Mth1p, *GAL80* was deleted from WT, *mth1*Δ, *std1*Δ and *mth1*Δ*std1*Δ strains. The deletion strains were grown in melibiose and galactose and the *P_GAL1_GFP* and *PMEL1mCherry* expression was monitored by microscopy **(Figure S4, S5, S6, S7)** as well as flowcytometry analysis **(Figure 6 and S8)**. Upon deletion of *GAL80* the *GAL* genes are constitutively expressed, and therefore the *P_GAL1_GFP* expression was observed when the cells were grown even in the non-inducible and non-repressible media i.e. gly/lac as the sole carbon source. The *GAL1* promoter was increased many folds but no significant change in the *MEL1* promoter expression was observed.

**Figure 6:**
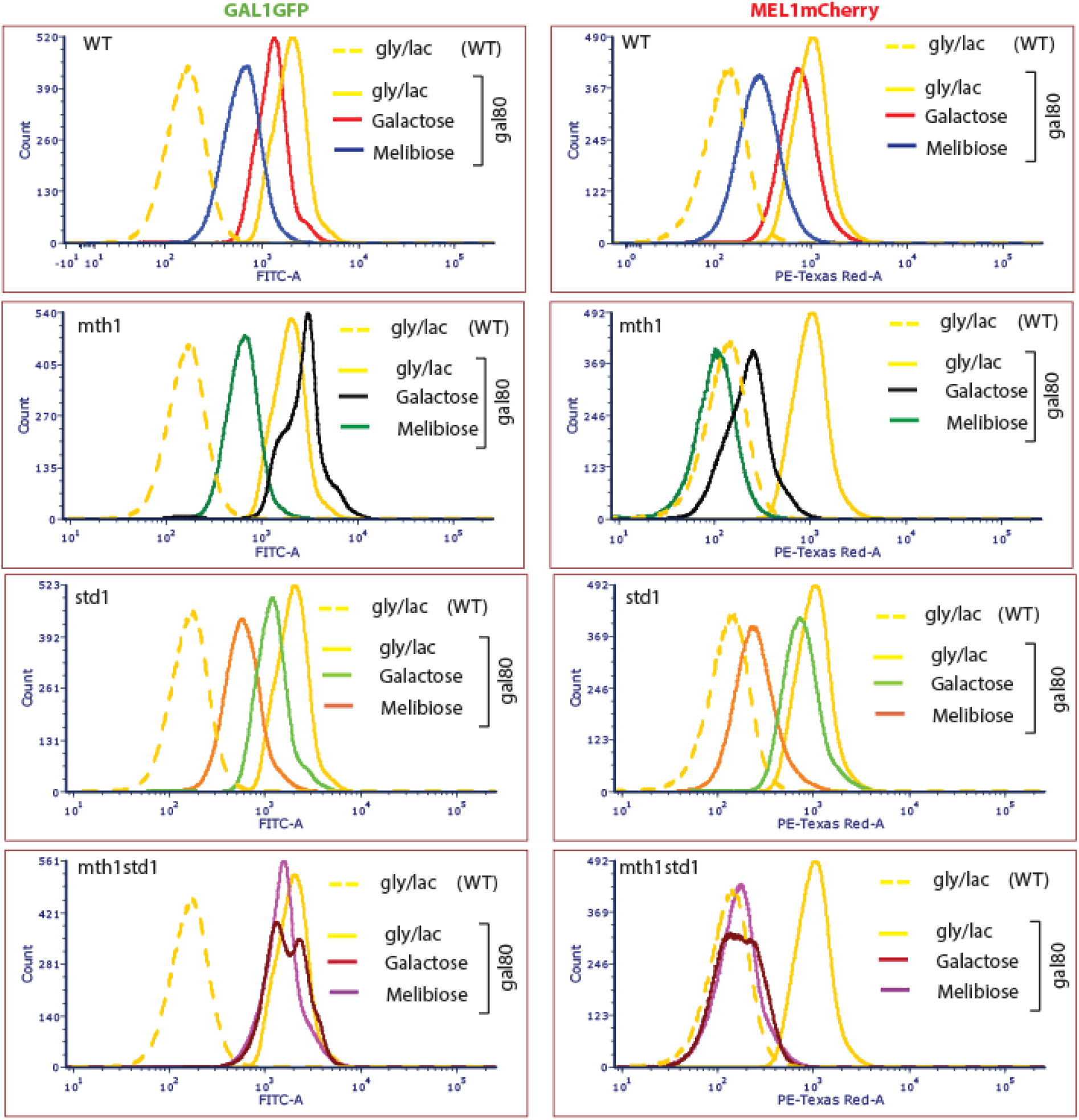
Flowcytometry analysis of the indicated strains integrated with *P_GAL1_GFP* and *P_MEL1_mCherry* grown in galactose and melibiose. WT, *mth1*Δ, *std1*Δ and *mth1*Δ*std1*Δ cells deleted for *GAL80* integrated with *P_GAL1_GFP* and *P_MEL1_mCherry* were monitored in flowcytometry after growing in the indicated media and mean fluorescence intensity was plotted. The experiment has been performed at least thrice and the pattern was same, therefore only one set of data is presented. Each data point represents the mean fluorescence intensity of 50,000 cells. GFP and m-Cherry were measured by using FITC and PE-Texas Red-A filters respectively.

## Discussion

Genome duplication and divergence establishes genetic novelty (Crow and Wagner, 2006; Hittinger, 2007). *Kluyveromyces lactis* and *Candida albicans*, are the ancestors of *Saccharomyces cerevisiae*, in which the genes have not undergone whole genome duplication and glucose signaling is regulated by a single gene, *SMS1* and *CaSTD1*, respectively. But in *S. cerevisiae*, after the whole genome duplication event, even though Mth1p and Std1p are dedicated to regulating the glucose signal transduction pathway, why only Mth1p is regulated by Gal4p and galactose mediated signaling is not been addressed so far. Therefore, we studied the role of this pair of paralogue genes, *MTH1* and *STD1*, that are the negative regulators of the glucose signal transduction pathway (Kaniak et al., 2004; Moriya and Johnston, 2004), but only one of them, i.e., *MTH1*, is under *GAL* switch regulation. The question then arises, after whole genome duplication, why are the *MTH1* and *STD1* regulation patterns assigned in an entirely different way. We surmised that, because Mth1p regulates the glucose signal transduction pathway and is positively regulated by galactose, it must have a role in regulating the utilization of complex disaccharides like melibiose, which is cleaved into glucose and galactose, that play a key role in repressing and inducing *MEL1* expression **(Figure S1B)**.

The phenotypic analysis by using 2-DG toxicity assay suggests that *GAL* genes are expressed in all cells pre-grown on galactose as the sole carbon source and are killed in 2-DG plates. But *GAL* genes are expressed only in a fraction of cells when pre-grown on melibiose as the sole carbon source, therefore only that fraction of cells (approximately 50%) are killed in 2-DG plates **(Figure 1)**. This data suggests that the melibiose grown cells show population heterogeneity i.e. one fraction of cells utilizes galactose and the other fraction utilizes glucose as a carbon source, which are cleaved from the disaccharide melibiose.

Based on our results, it can be inferred that even melibiose, which hydrolyzes into glucose and galactose, is utilized in a different strategy by the cells than the equimolar mixture of glucose and galactose **(Figure 1 and 2)**. Melibiose-utilizing cells exhibit cellular heterogeneity among themselves, and a fraction of cells utilizes glucose and another fraction utilizes galactose. The glucose liberated from the melibiose hydrolysis seems to be not above the threshold level to repress the *GAL* genes of the cells, which are utilizing the galactose from melibiose hydrolysis. This shows the division of labor among the same population to utilize carbon sources efficiently in an fluctuating environment (Kavatalkar et al., 2020). When the cells were grown on an equimolar mixture of glucose and galactose, no expression of *GAL* genes in the log phase was observed. However, at the stationary phase when glucose is completely utilized, *GAL* genes expression was observed. That is, when a glucose and galactose mixture is utilized as a carbon source, the diauxic shift is observed but when melibiose is utilized as the sole carbon source, no diauxic shift was observed **(Figure 2A)**. This suggests that at log phase the cells preferentially utilize glucose as a carbon source and once glucose is completely exhausted from the medium galactose utilization takes place. This is becoming a possibility because during growth on melibiose, the amount of α-galactosidase expressed is only enough to produce very low concentration of galactose and glucose and *MTH1* seems to play a role in regulating the α-galactosidase expression.

It is also difficult to appreciate the functional history of *MTH1* and *STD1* in melibiose utilization without comparing it with its homologue *SMS1* of *K. lactis* (Hnatova et al., 2008; Wolfe and Shields, 1997). In the ancestors *K. lactis* and *C. albicans*, the glucose signaling is regulated by a single gene *SMS1* and *CaSTD1*, respectively. But in *S. cerevisiae* it is regulated by a duplicate pair of genes *MTH1* and *STD1*, and *MTH1* is also induced by galactose. Based on our results, it is observed that upon deletion of *SMS1* in *K. lactis*, growth on lactose is not hampered **(Figure 3A)**like the deletion of either *MTH1* or deletion of *MTH1* and *STD1* together in a WT *S. cerevisiae* strain **(Figure 3B)**. Therefore, we suggest that the duplication and divergence of *SMS1* of *K. lactis* to *MTH1* and *STD1*, gave ample opportunities to *Saccharomyces* to become an efficient disaccharide user. It has been shown that, *S. cerevisiae* has altered the GAL switch as compared to some ancestors to lose the ultra-sensitivity and stringent glucose repression. These changes caused an increase in fitness in the disaccharide melibiose at the expense of a decrease in fitness in galactose (Das Adhikari et al., 2014). But the deletion of *MTH1* (*mth1*Δ) shows the growth defect **(Figure 3C** and **D)** as well as reduction of only the *MEL1* promoter-driven *mCherry* expression, but not the *GAL1* promoter-driven *GFP* expression in either galactose or melibiose as the sole carbon source. However, these defects were not observed on the deletion of its paralogue *STD1* (*std1*Δ). These defects in the growth, as well as expression, were found to be enhanced upon deletion of both *MTH1* and its paralogue *STD1* (*mth1*Δ*std1*Δ) together. The *mth1*Δ strain not only shows the growth defect, but also shows a marginal reduction in biomass production compared to the WT and *std1*Δ strain because of non-efficient carbon source utilization **(Figure 3C and D)**. This cell-to-cell heterogeneity in melibiose-grown cells is corroborated by the microscopic and flowcytometry analysis in *P_GAL1_GFP* and *P_MEL1_mCherry* integrated cells, where some very bright *P_GAL1_GFP* cells are clearly visible in the merged panel in melibiose grown cells indicated with white arrows **(Figure 4 and 5)**.

Another interesting observation of our study is that even though both the *GAL1* and *MEL1* promoter are regulated by same galactose signal driven Gal3p mediated Gal4p-Gal80p interaction mechanism, we found that Mth1p regulates only the *MEL1*, which has single Gal4p binding sites, but not *GAL1*, which has multiple Gal4p binding sites, by some unknown interaction other than the galactose signal driven Gal3p mediated Gal4p-Gal80p interaction mechanism **(Figure 4 and 5 and S3).**This is because upon deletion of *mth1* or *mth1std1* the *P_GAL1_GFP* expression is not affected in any panel but *P_MEL1_mCherry* is almost nil in *mth1*Δ or *mth1*Δ*std1*Δ **(Figure 4 and 5 and S3)**.

This suggests that *MTH1* plays a key role in the utilization of the disaccharide melibiose, which is cleaved by α-galactosidase into equimolar concentrations of glucose and galactose, while *STD1* may not have that important role with respect to melibiose utilization.

Our inference from this study rules out the possibility of Mth1p-mediated inhibition of the Gal80p-Gal4p interaction in the Gal3p mediated galactose signal transduction of the *GAL1* promoter driven *GAL* genetic switch. Therefore, surprisingly upon the deletion of *GAL80* in the WT, *mth1*Δ, *std1*Δ or *mth1*Δ*std1*Δ strains, only the *GAL1* promoter driven GFP expression was increased, but no significant change in the *MEL1* promoter driven mCherry expression was observed, in the presence of either galactose or melibiose **(Figure 6 and S8)**.

We also infer that *MEL1* has a basal and low expression, therefore no clear bimodal population can be seen in the MEL1mcherry panel **(Fig 5D and 6D)**. But in the GAL1GFP panel bimodal population is observed because the *GAL1* promoter is tightly regulated and has no basal expression, so two populations coexist, one in which *GAL* genes are not expressed, and in the other population *GAL* genes are expressed **(Fig 5D and 6D)**. But, when the *GAL80* is deleted that uninduced population is also getting constitutively expressed and still both the 2 populations coexist but their *GAL* gene expression level is different and still bimodal population coexists **(Fig 5D and 6D)**. Together both *MTH1* and *STD1* repress the glucose effect well rather than the *MTH1* alone. That is why double deletion *mth1*Δ*std1*Δ has a more prominent effect and clear bimodality is observed **(Fig 5D and 6D)**.

This suggests that Mth1p might play a role on the downstream of the Gal80p-Gal4p interaction in the galactose signal transduction in the *GAL* genetic switch, which suggests that Mth1p might be regulating the *MEL1* promoter beyond Gal4p and Gal80p interaction by some different mechanism.

Based on our results and with the available evidence, we propose a model of the role of *MTH1* in *GAL* switch regulation **(Figure 7)**. The mechanism for utilization of melibiose is like the “chicken and egg” story, i.e., whether the α-galactosidase is encoded first, or the galactose is generated first from the hydrolysis of melibiose to induce α-galactosidase generation, is not clearly understood. Therefore, understanding the mechanism of melibiose utilization will clarify our understanding of the mechanism of other complex sugars utilization, like sucrose and raffinose, but also the reversible conversion of ethanol to acetaldehyde by *ADH1* and *ADH2* genes in the yeast (de Smidt et al., 2008), which are controlled by feedback. This sort of understanding may have potential application on biotechnological applications like amino acid overproduction (Lee et al., 2018; Qin et al., 2015), Sugar ethanol (Ostergaard et al., 2000; Oud et al., 2012; Rønnow et al., 1999), biofuel, and Dairy industries.

**Figure 7:**
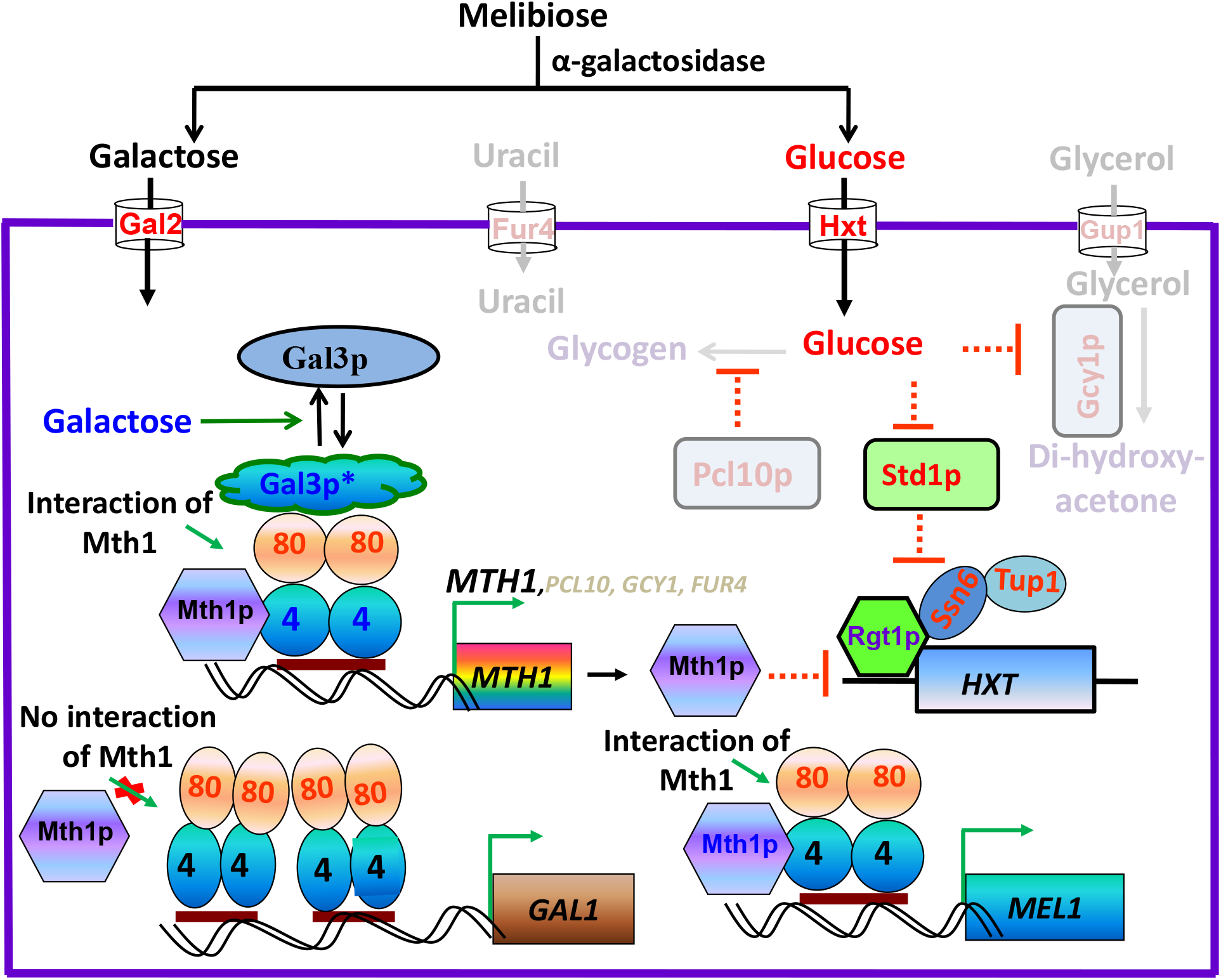
A proposed model to show the comparison of mode of action of Mth1p in single and multiple Gal4p binding sites in *GAL* and *MEL* promoter. Mth1p along with Rgt1p represses the hexose transporters that transport glucose into the cell, which is generated by cleaving of melibiose. Mth1p seems to activate the single Gal4p binding sites like in *MEL1* gene, but not the double Gal4p binding sites like in *GAL1* gene. *MTH1*, *PCL10*, *GCY1* and *FUR4* are regulated by Gal4p-Gal80p binding and induced by galactose, but only *MTH1* is repressed by glucose, therefore *PCL10, GCY1* and *FUR4* are shown in light color. Gal3p* is the active Gal3 protein. Green arrows represent activation and the red blunt ended lines represent the repression.

The limitation to our study is the difficulty in quantifying the glucose, galactose and melibiose in the spent media, as many other constituents are there in the synthetic media and it is hard to quantify galactose and melibiose, as melibiose also contains the galactose moiety.

In the concluding remarks, in this communication we present given experimental evidence on how the disaccharide, melibiose, is utilized by yeast due to population heterogeneity in the cells and the role of *MTH1* gene in balancing the tuning of glucose and galactose upon hydrolysis of the disaccharide melibiose.

## Materials and methods

### Yeast strains, media and growth conditions

Yeast strains **(Table S1)**were grown either in the yeast extract, peptone and dextrose (YEPD) or in the synthetic medium containing 3% (v/v) glycerol plus 2% (v/v) potassium lactate (gly/lac), glucose or galactose (Sigma) as the carbon source. Yeast transformation was done by the lithium acetate protocol (Gietz and Woods, 2002). Transformation of *E. coli*, plasmid amplification and isolation was carried out by standard protocols (Sambrook, 1989).

#### Plasmids construction

##### Construction of *pGALlGFP*

A 1.5 kb cassette containing *P_GAL1_GFP* isolated from *YCpGAL1-GFP*(Stagoj et al., 2005) as a *Pst*I-*Eco*RI fragment was ligated at the corresponding sites of *YIpLac211* to obtain *pGAL1-GFP. pGAL1-GFP* was linearized with *EcoRV* located within the *URA3* and integrated in the strains (Kar et al., 2017; Kar et al., 2014).

##### Construction of *pRKK3*

*MEL1* promoter was amplified from genomic DNA by primer PJB 415 (5’ CACAGGAAA CAGCTATGACCATGATTACGCCAAGCTTGCATGCCTGCAGGATTCTTTCTGTACGCTCAGGGTGGGC3’) and PJB 416 (5’TGCTCCAGCACCAGCACCAGCACCTGCTCCGGATCCTCTAGAGTCGACGTCGTTGCTTTTATTACCGTTGCTC 3’) and cloned to *YEpLac195-DsRed-Kanamycin* plasmid cut with *Bam*HI by the in-vivo cloning method. Then the DsRed-Kanamycin fragment was removed from the *YEpLac195-DsRed-Kanamycin* by BamHI and *Eco*RI. The *mCherry-Kanamycin* fragment was PCR amplified from plasmid *pWA8-mCherry* (Euroscarff) by PJB435 (5’ CATGTCGCGGATCCATGGTGAGCAAGGGCGAGGAGGATA 3’) and PJB436 (5’ CGTTGTAAAACGACGGCCAGTGAATTCGCGCGGCCGCATAACTTC G 3’). The PCR product was digested with *BamHI* and *Eco*RI and ligated to the *BamHI* and *Eco*RI digested *YEpLac195-DsRed-Kanamycin* resulting a *YEpLac195-MEL1-mCherry-Kanamycin* clone **(Table S2)** (Das Adhikari and Bhat, 2016).

##### Construction of *pRKK4*

The *pRKK4* (*YEpLac195-MEL1-m-CherryKan*) was digested by the *Sa*lI restriction enzyme, which gives one fragment containing *PMEL1mCherry-Kan.This* fragment was cloned to the Trp locus containing Integrative vector *YIpLac204* at *Sa*lI site of the multi cloning site and screened by blue white screening **(Table S2)**.

#### Strain construction

##### Disruption of *MTH1, STD1* and *GAL80* genes

###### *MTH1* disruption

*MTH1* was disrupted in different strain backgrounds by integrating the *mth1:KanMX4* cassette. This cassette was obtained by amplifying genomic DNA of *BY4742*, which contains *mth1:Kanamycin* cassette using primers PJB419 (5’CATCGTGAGAGAAAATACGAGTCCATTTCTCCAGTGAAACTACCGTAG3’) and PJB420 (5’GACATTTACCGCTTGCGCGCGGCTTCTTCTGTATGCTGTGTC3’). The resulting PCR fragment was flanking *KanMX4* cassette by 389 bp of upstream sequence of ATG and 372 bp from downstream of start and stop codon of *MTH1* ORF. The strains needing to be disrupted were transformed with this PCR product and the transformants were selected in YEPD plates supplemented with G418 to a final concentration of 200μg/ml. The *mth1*Δ was confirmed by PCR amplification using primers PJB426 (5’GAAGACTCTTGAGGAGGTAGGG3’) 881 bp upstream of start codon of *MTH1* ORF and PJB427 (5’CTCTGGCGCATCGGGCTTCCC3’) 605 bp downstream of start codon of *Kanamycin* ORF.

###### *STD1* disruption

*STD1* was disrupted in different strain backgrounds by integrating the *std1:Hphr* cassette. This cassette was obtained by amplifying genomic DNA of KFY917 strain, which contains *std1:Hphr* cassette using primers PJB635 (5’GGGAAGTTCATGCTATACAACGCC3’) and PJB636 (5’GCTAGTAAATC GGCCGGATCTC3’). The resulting PCR fragment was flanking *Hphr* cassette by 495 bp of upstream sequence of ATG and 474 bp from downstream of start and stop codon of *STD1* ORF. The strains needing to be disrupted were transformed with this PCR product and the transformants were selected in a YEPD plate supplemented with hygromycin to a final concentration of 100μg/ml. The *std1*Δ was confirmed by PCR amplification using primers PJB637 (5’CGGAATAACAAGCAAAGTCGG3’) 999 bp upstream of start codon of *STD1* ORF and PJB471 (5’CCCCGAACATCGCCTCGCTCC3’) 679 bp downstream of start codon of *Hygromycin* ORF (*Hphr*).

###### *GAL80* disruption

*GAL80* was disrupted in the strains by an integrating *gal80:LEU2* cassette. This cassette was obtained by amplifying genomic DNA of BY2685 strain, which contains *gal80:LEU2* cassette using primers PJB485 (5’CCTCCTCCAGATGGAATCCCTTCCATAG3’) and PJB486 (5’GCAAACCTATC ACCCGGTGATAACAGC3’). The resulting PCR fragment was flanking *LEU2* by 507 bp of upstream sequence of ATG and 479 bp from downstream of start and stop codon of *GAL80* ORF. The strains needed to be disrupted were transformed with this PCR product and transformants were selected in leucine drop out plate. The *gal80*Δ was confirmed by phenotypic analysis and by PCR amplification using primers PJB544 (5’GCCTGTCTACAGGATAAAGAC3’) 1000 bp upstream of start codon of *GAL80* ORF and PJB27 (5’CGCCAAGCTTGATATGAG3’) 20 bp upstream of stop codon of *GAL80* ORF.

##### Integration of *pGAL1GFP* and *pRKK4* into WT, *mth1*Δ, *std1*Δ, *mth1*Δ*std1*Δ and their respective *gal80*Δ stains

The *pGAL1GFP* and *pRKK4* were digested with *Eco*RV enzyme and integrated at the *URA3* and *TRP1* locus of the WT, *mth1*Δ, *std1*Δ, *mth1*Δ*std1*Δ and their respective *gal80*Δ strains to obtain the respective integrant strains **(Table S1)**. The *MTH1, STD1* and *GAL80* were deleted in the strains as mentioned above and their respective deletion strains were obtained **(Table S1)**. After the integration or the deletion, the strains were characterized either by their fluorescence expression or by confirmation PCR.

### Growth curve analysis

Cells were pre-grown on gly/lac and again inoculated onto different concentrations of filter-sterilized melibiose or galactose or glucose or glucose and galactose mixture. The OD was monitored at 600nm, at an interval of 2 hours and a few time points at 4 or 6 hours by taking out a small volume of cells from the 50 ml flask culture. The OD was plotted against time and the growth pattern was analyzed from the obtained curve.

### 2-Deoxy galactose (2-DG) toxicity assay

Constitutive expressions of *GAL1* in various medium-grown cells were monitored based on their ability to grow in gly/lac + 2-DG (0.3%). It has been reported that cells that are constitutive for galactokinase expression would not be able to grow in a media containing 2-DG due to toxicity (Platt 1984). This toxicity is due to the accumulation of the 2-deoxygalactose-1-phosphate (2-DG-1-p), which cannot be further metabolized.

### Microscopic analysis

Cells were grown in complete medium containing gly/lac and re-inoculated into medium containing 2% glucose and 2% of galactose. Cells were harvested, washed with PBS twice, re-suspended in PBS and kept in ice. The images were acquired using an Olympus microscope with FluoView^TM^ application software equipped with a 100x objective. The EGFP and Texas red filters were used to detect the expression of *P_GAL1_GFP* and *PMEL1mCherry*, respectively.

### Flow-cytometry analysis (FACS)

For flowcytometry analysis, the samples were prepared using the same procedure as for microscopic samples. A minimum of 50,000 cells were analyzed for each data point. Flow-cytometry data were collected using BD FACS ARIA-I flow-cytometer with FITC and Texas red filters for detection of the expression of *P_GAL1_GFP* and *P_MEL1_mCherry*, respectively. Data were collected by BDFACS DIVA software. All experiments were performed at least thrice, and the patterns were found to be same. Therefore, only one set of data is compared and presented here.

## Abbreviation

Gal3p: Gal3 protein.
Δ: deletion
WT: wild type

## Acknowledgements

We thank CRNTS and SAIF of IITB for confocal microscopy and flowcytometry facility. We acknowledge the financial support from IITB, CSIR to RKK and, DST and 08BRNS004 to PJB. We thank Dr. Komel, Dr C. Wittenberg and Dr M. Lemaire for generously gifting me strains and plasmids. We also thanks PJB lab members and Dr. Edith Wolf of NIDCR, National institute of critical suggestion on manuscript.

## Author Contributions

Conceived by PJB and designed the experiments by: RKK and PJB. Performed the experiments: RKK. Analyzed the data: RKK and PJB. Contributed reagents/materials/analysis tools: RKK and PJB. Wrote the paper: RKK.

## Competing Interest

The authors have declared no competing interest.

