## Supporting information for "Cellular heterogeneity and *MTH1* play key roles in galactose mediated signaling of the GAL switch to utilize the disaccharide melibiose"

Fax: + (91-22) 2572 3480

Phone: + 16086092552

### Supplementary figures

**Supplementary figure S1:**

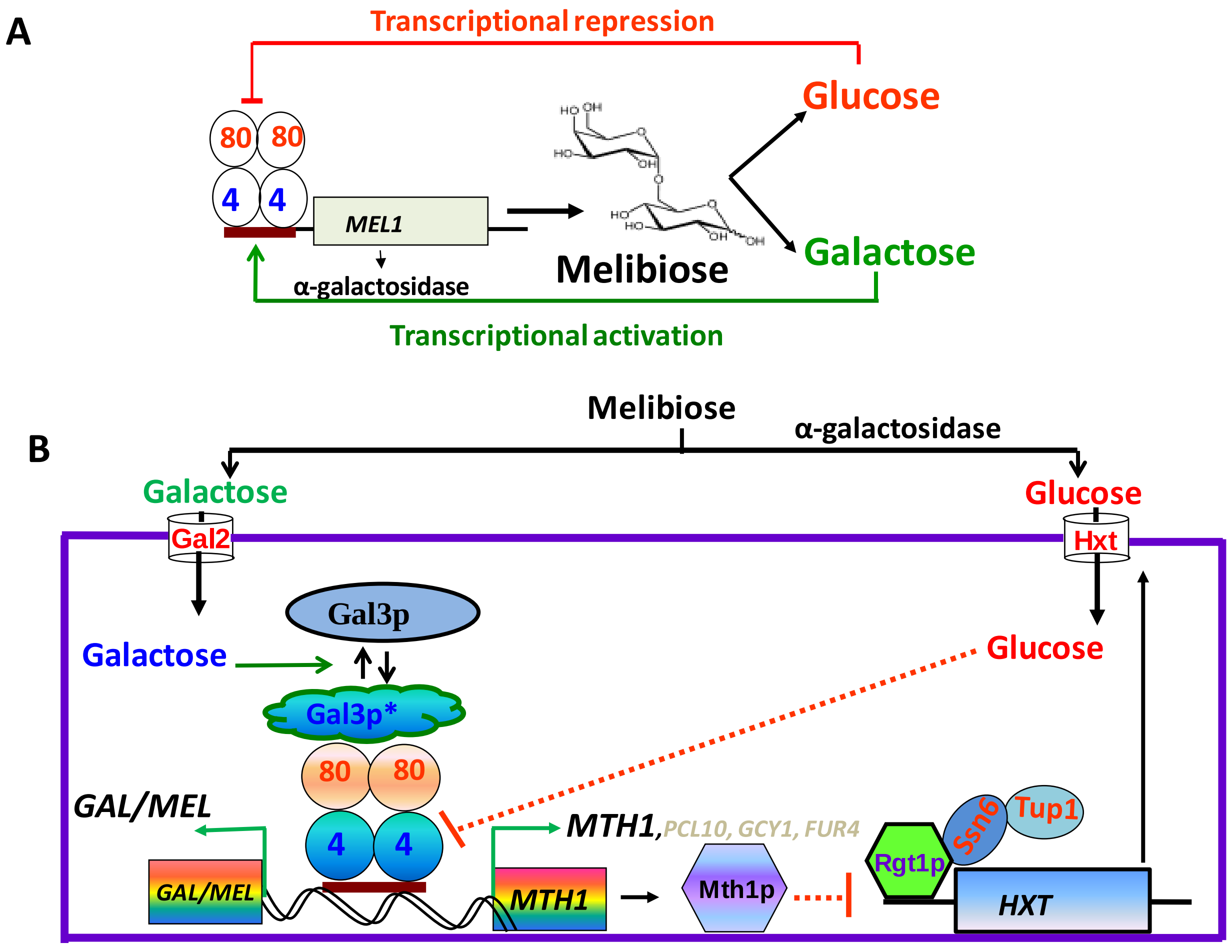


**Supplementary figure S1. Model shows the regulation of *MEL1* and role of *MTH1* in glucose, galactose signal regulation**. **(A)** Melibiose is cleaved into equimolar concentration of glucose and galactose. Glucose represses the *MEL1* expression (red line with the blunt end), and galactose induces the activation of *MEL1* expression (green line with the arrow). **(B)** *MTH1* is regulated by Gal4p-Gal80p (**4-80**) interaction, same as the *MEL1* gene or other *GAL* genes. *MTH1* is repressed by glucose, induced by galactose that is generated upon melibiose hydrolysis, and repressed the *HXT* genes that transport glucose. *PCL10*, *GCY1* and *FUR4* are induced by galactose but not repressed by glucose, therefore are written in light color.

**Supplementary figure S2:**

A

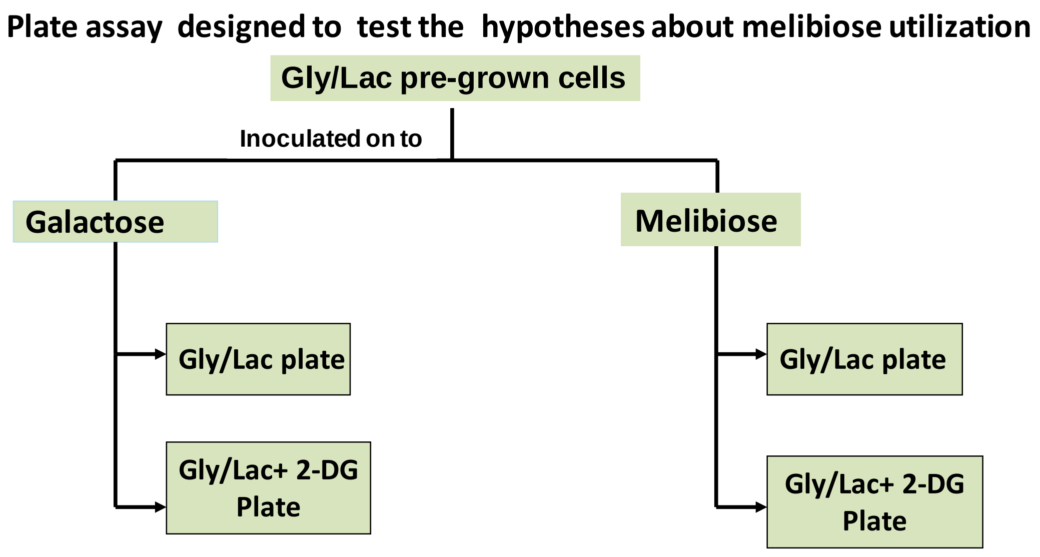


**Supplementary figure S2. (A)** Schematic of the 2-Deoxy Galactose plate assay to score the number of cells that responds to galactose, when they are pre-grown in melibiose or galactose.

**Supplementary figure S3:**

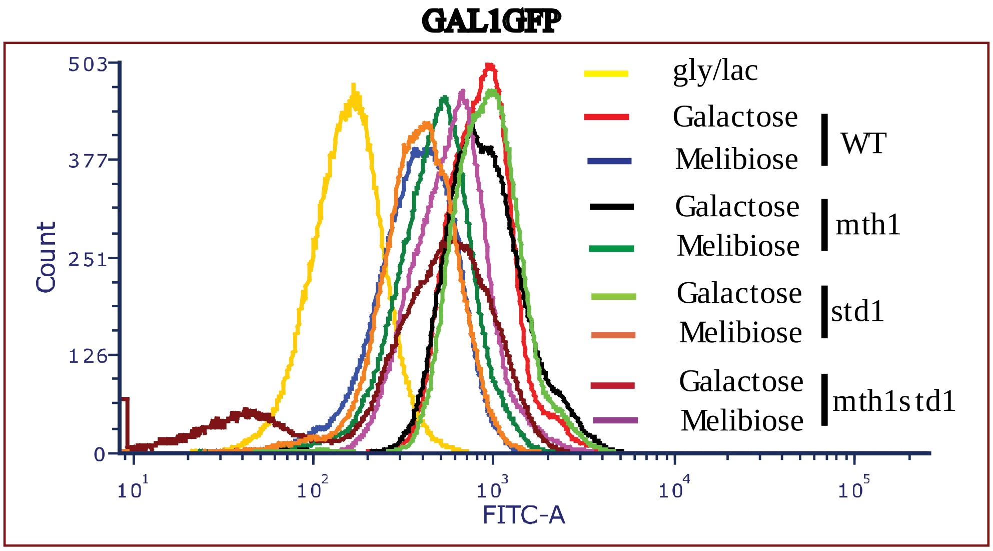


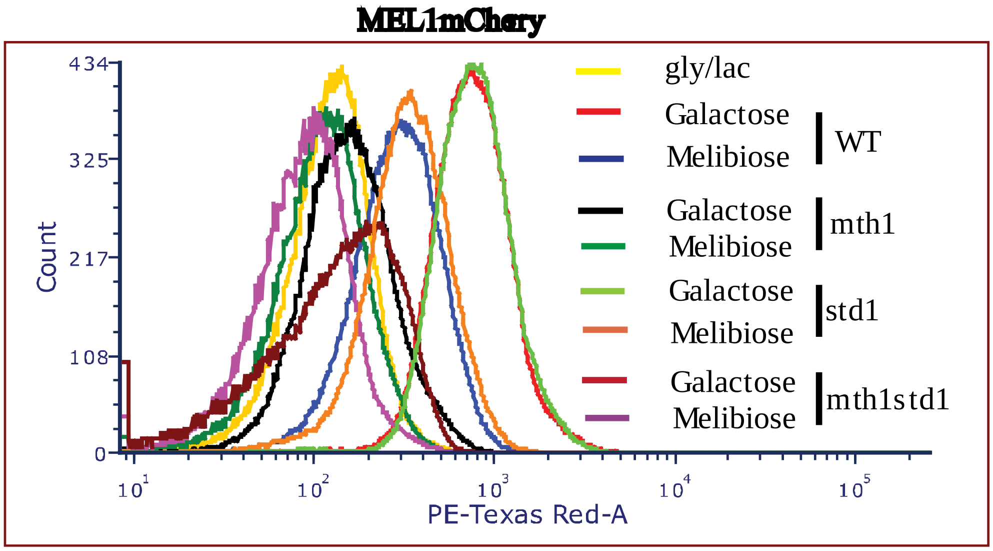


**Figure S3: Flowcytometry analysis of the indicated strains integrated with *$P_{GAL1}$GFP* and *$P_{MEL1}$mCherry* grown in galactose and melibiose**. WT, *mth1*Δ, *std1*Δ and *mth1*Δ*std1*Δ cells were integrated with *$P_{GAL1}$GFP* and *$P_{MEL1}$mCherry* were monitored in flowcytometry after growing in the indicated media and mean fluorescence intensity was plotted. The experiment has been performed at least thrice and the pattern was same, therefore only one set of data is presented. Each data point represents the mean fluorescence intensity of 50,000 cells. GFP and m-Cherry were measured by using FITC and PE-Texas Red-A filters respectively.

**Supplementary figure S4:**

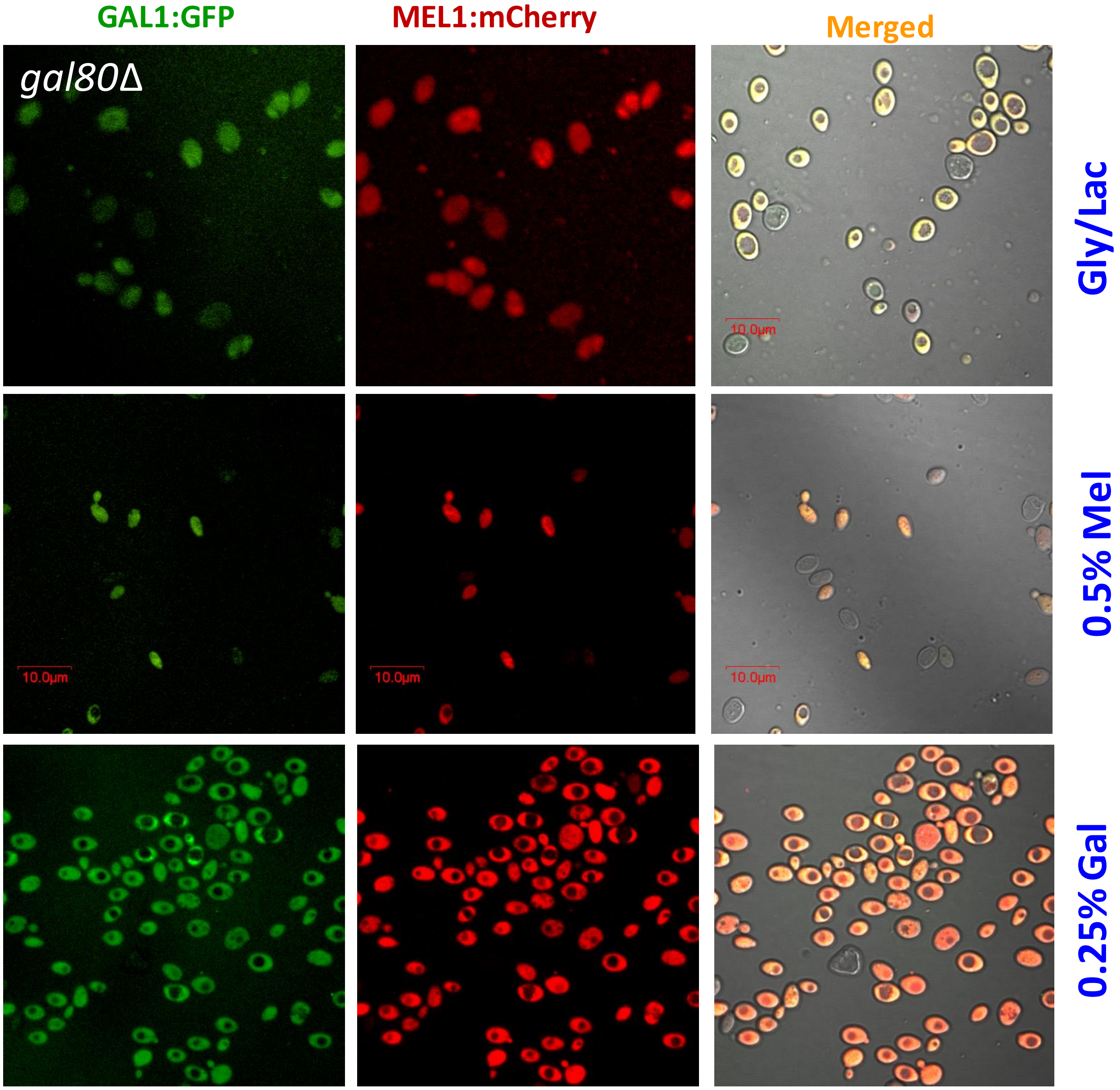


**Supplementary figure S4.** **Microscopic analysis of *gal80Δ* strains integrated with *$P_{GAL1}$GFP* and *$P_{MEL1}$mCherry* grown in galactose and melibiose**. *gal80Δ* strain integrated with *$P_{GAL1}$GFP* and *$P_{MEL1}$mCherey* were grown on the indicated media and monitored in a confocal microscope. The scale bar is 10 μm. GFP and m-Cherry were monitored by using EGFP and PE-Texas Red filters respectively.

**Supplementary figure S5:**

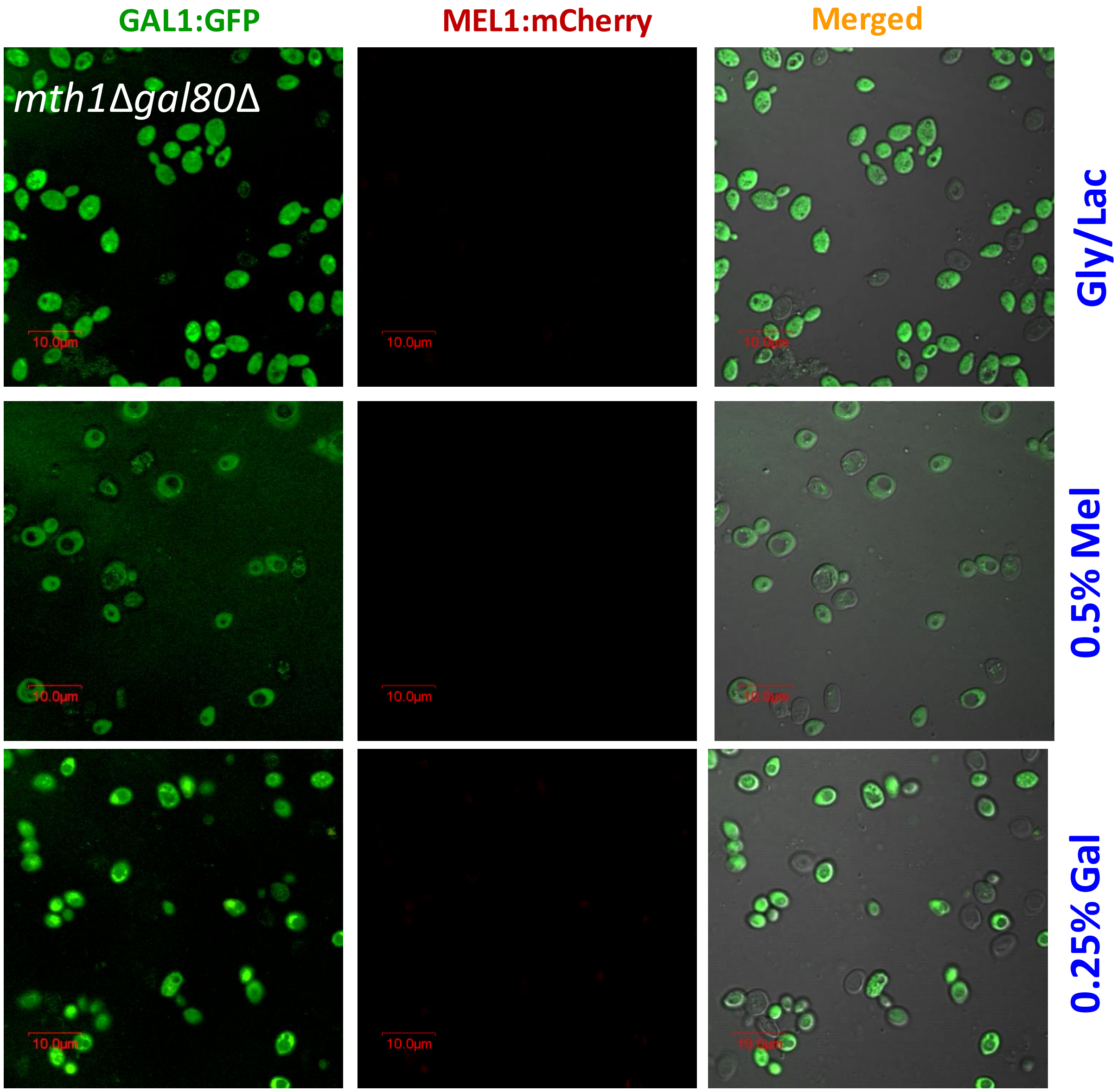


**Supplementary figure S5.** Microscopic analysis of ***mth1Δgal80Δ* strains integrated with *$P_{GAL1}$GFP* and *$P_{MEL1}$mCherry* grown in galactose and melibiose**. Other conditions are the same as figure S4.

**Supplementary figure S6:**

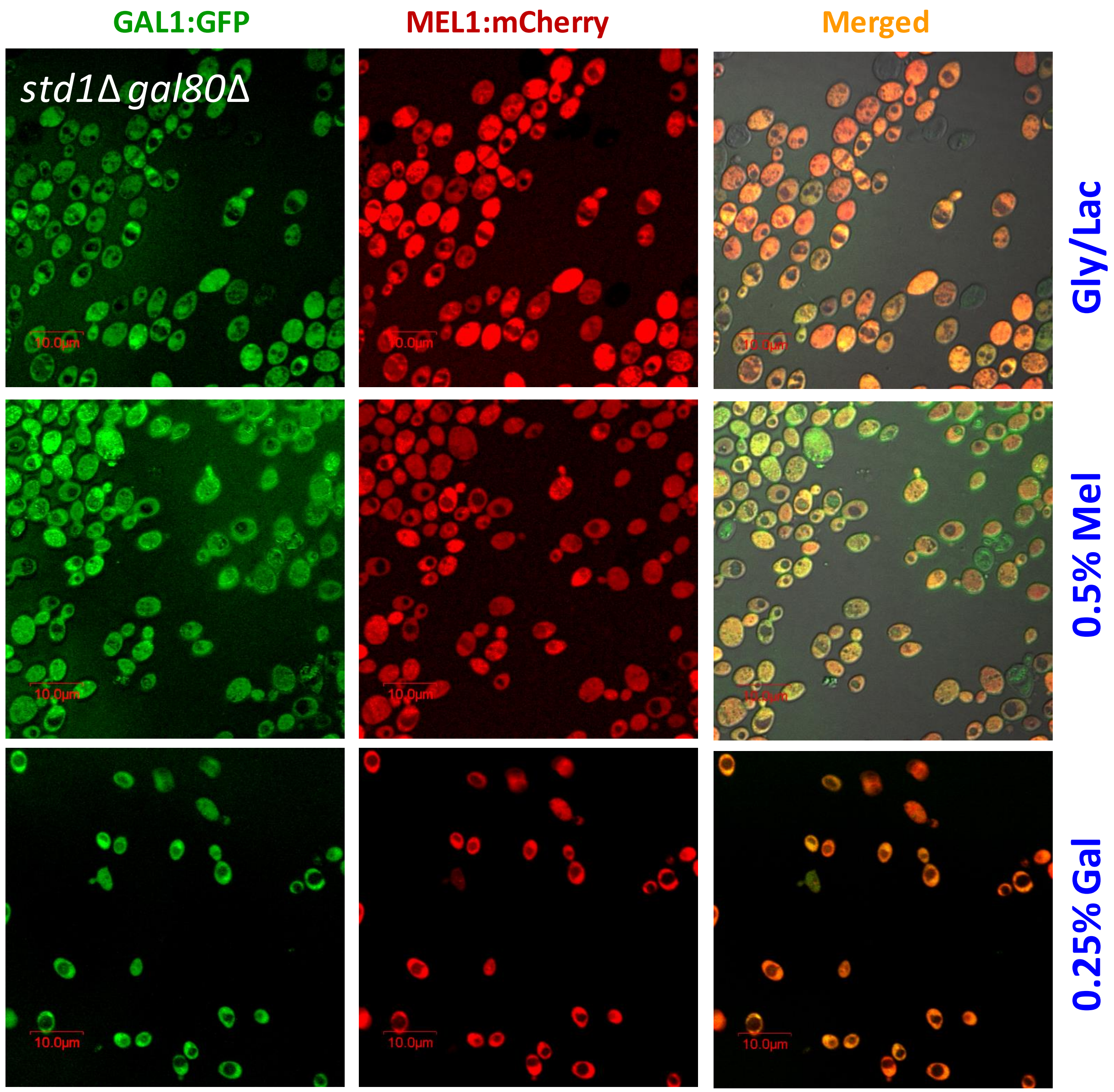


**Supplementary figure S6.** Microscopic analysis of ***std1Δgal80Δ* strains integrated with *P<sub>GAL1</sub>GFP* and *P<sub>MEL1</sub>mCherry* grown in galactose and melibiose**. Other conditions are the same as figure S4.

**Supplementary figure S7:**

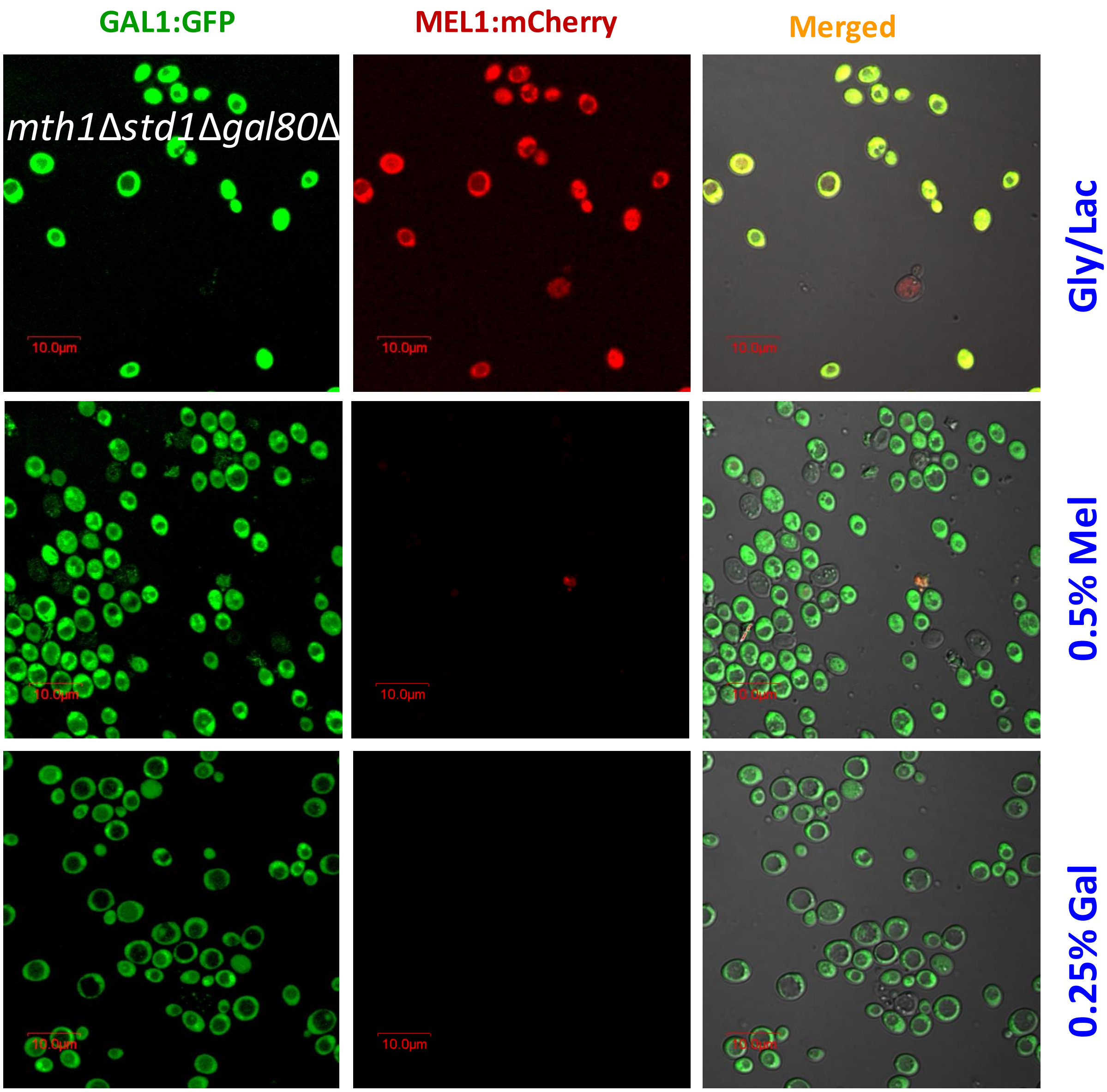


**Supplementary figure S7.** Microscopic analysis of ***mth1*Δ*std1*Δ *gal80*Δ strains integrated with *P*<sub></sub>** **$P_{GAL1}$*GFP* and $P_{MEL1}$*mCherry* grown in galactose and melibiose**. Other conditions are the same as figure S4.

**Supplementary figure S8:**

GAL1GFP

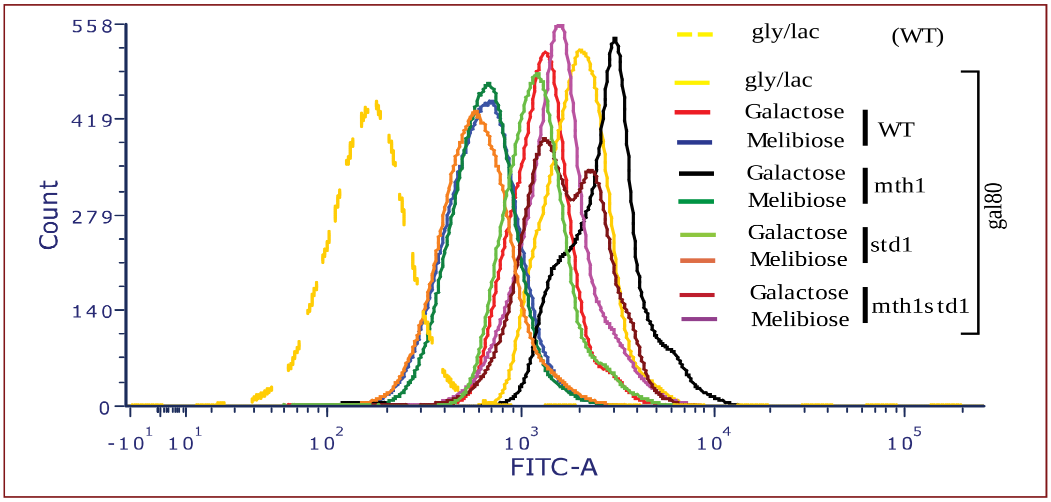


MEL1mCherry

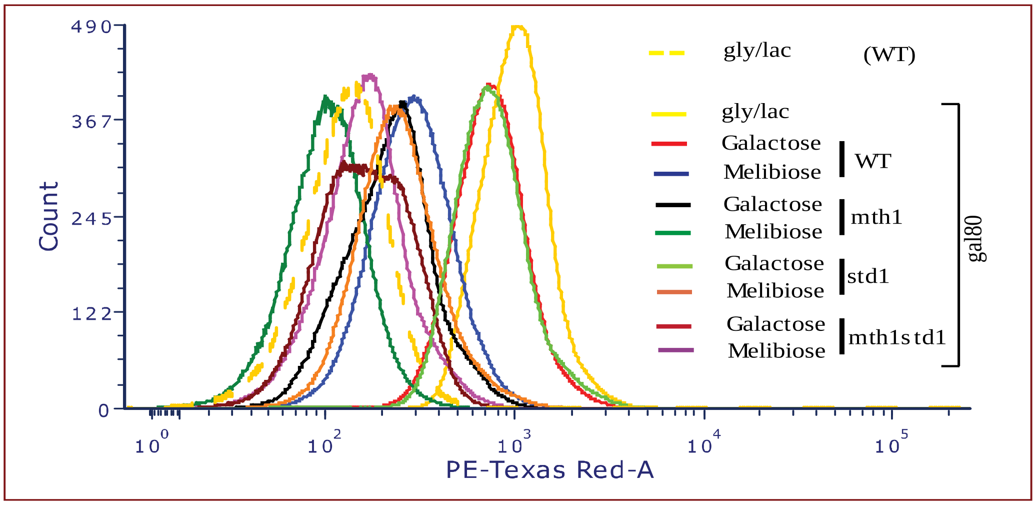


**Figure S8: Flowcytometry analysis of the indicated strains integrated with $P_{GAL1}$*GFP* and $P_{MEL1}$*mCherry* grown in galactose and melibiose**. WT, *mth1*Δ*, std1*Δ and *mth1*Δ*std1*Δ cells deleted for *GAL80* integrated with $P_{GAL1}$*GFP* and $P_{MEL1}$*mCherry* were monitored in flowcytometry after growing in the indicated media and mean fluorescence intensity was plotted. The experiment has been performed at least thrice and the pattern was same, therefore only one set of data is presented. Each data point represents the mean fluorescence intensity of 50,000 cells. GFP and m-Cherry were measured by using FITC and PE-Texas Red-A filters respectively.

**Supplementary Table S1: List of strains used in this study and their genotypes.**

| Strain | Genotype | Source |
|---|---|---|
| *MW270-7B* | *MATa uraA1-1 leu2 metA1-1* | Hnatova et al |
| MLK54 | *MATa uraA1-1 leu2 metA1-1:sms1::kanMX4* | Hnatova et al |
| Sc723 | *MATα ade1 ile leu* 2-3, 112 *ura* 3-52 *trp1*-HIII *his3*- Δ1 *MEL1 lys2*::$GAL1_{UAS}$-$GAL1_{TATA}$-*HIS*3 | Hopper J |
| BY4742 | *MATα , his3Δ 1, leu2Δ 0, lys2Δ 0; ura3Δ 0 ,mth1*::*KanMx4* | Euroscarf |
| BY2685 | *MATa leu2 ura3 trp1 his3 ade gal80::LEU2* | YGRC , Japan |
| KFY917 | *MATa grr1LEU2 mth1ZEOstd1HphMX4 RGT1-3HAKanMX2* | Wittenberg |
| ScRKK7 | as Sc723 , but *URA3::* $P_{GAL1}$*GFP* | This study |
| ScRKK19 | as Sc723 , but *mth1*::*KanMx4* | This study |
| ScRKK20 | as ScRKK19, but *URA3::* $P_{GAL1}$*GFP* | This study |
| ScRKK21 | as ScRKK7, but *TRP2::* $P_{MEL1}$*mCherry* | This study |
| ScRKK22 | as Sc723, but *TRP2::* $P_{MEL1}$*mCherry* | This study |
| ScRKK23 | as ScRKK20, but *TRP2::* $P_{MEL1}$*mCherry* | This study |
| ScRKK24 | as Sc723 , but *std1*::$Hph^r$ | This study |
| ScRKK25 | as ScRKK19, but *std1*::$Hph^r$ | This study |
| ScRKK26 | as ScRKK21, but *std1*::$Hph^r$ | This study |
| ScRKK27 | as ScRKK23, but *std1*::$Hph^r$ | This study |
| ScRKK28 | as ScRKK21, but *gal80*::*LEU2* | This study |
| ScRKK29 | as ScRKK22, but *gal80*::*LEU2* | This study |
| ScRKK30 | as ScRKK26, but *gal80*::*LEU2* | This study |
| ScRKK31 | as ScRKK27, but *gal80*::*LEU2* | This study |

**Supplementary Table S2: List of plasmids**

| Plasmid | Features | Source |
|---|---|---|
| *YIpLac211Gal1GFP* | Integrative plasmid with *ura3*::*URA3GAL1GFP*. | Lab Stock |
| *pFA6KanMX4* | Kanamycin based plasmid | Euroscarff |
| *pRKK1* *YEpLac195-DsRedKan* | Multicopy plasmid containing *DsRedKan* cassette in YEpLac195 | This study |
| *pRKK2* *YEpLac195-MEL1:DsRedKan* | Multicopy plasmid containing *MEL1DsRedKan* cassette in YEpLac195 | This study |
| *YIpLac204* | Trp based Integrative plasmid | Lab stock |
| *pRKK3* *YEpLac195MEL1:mCherryKan* | Multicopy plasmid containing *MEL1*:mCherryKan cassette in YEpLac195 | This study |
| *pRKK4* *YIpLac204MEL1:mCherryKan* | Integrative plasmid containing *MEL1*:mCherryKan cassette in YIpLac204 | This study |
| *pAW8mCherry* | mCherry with Kanamycin marker | Euroscarff |
